# A Comparative Assessment of *edgeR* and *methylKit* Pipelines for DNA Methylation Detection

**DOI:** 10.1101/2025.05.11.653026

**Authors:** Iraia Muñoa-Hoyos, Manu Araolaza, Irune Calzado, Mikel Albizuri, Nerea Subirán

## Abstract

Despite the improvements in tool development for DNA methylation analysis, there is a lack of a consensus on computational and statistical models used for differentially methylated cytosine (DMC) identification. This variability complicates the interpretation of findings and raises concerns about the reproducibility and biological significance of the detected results. In this regard, the primary objective of this study was to compare the performance, concordance, and biological relevance of *edgeR* and *methylKit* tools in detecting DMCs (the first one based on fold change and the second one based on percentage), following morphine exposure model in mouse embryonic stem cells (mESCs). While a different number of total DMCs was identified by each tool, both pipelines detected a global hypomethylation as a result. Genomic analysis revealed a predominant distribution of DMCs in intergenic and intronic regions on one hand, and in open sea regions on the other hand. Despite the differences in sensitivity, both tools demonstrated moderate concordance in DMCs detection (∼56%) and high concordance in gene level analysis (∼90%), identifying similar differentially methylated genes (DMGs). Overall, the results underscore the complementary strengths of *methylKit* and *edgeR* and highlight the importance of tool selection for epigenetic studies. As a conclusion, integrating both pipelines is recommended for comprehensive analysis, particularly in studies with complex experimental designs.

## INTRODUCTION

Epigenetics is the study of heritable changes in gene expression that do not involve alterations to the underlying DNA sequence. The primary mechanisms of epigenetic regulation include DNA methylation, histone modifications, chromatin remodelling and RNA based regulation (involving both small and long non-coding RNA). These modifications are essential for cellular differentiation and function, as well as for normal development, aging and a number of diseases (Moosavi A et al. 2016; Evangelina R et al. 2025).

Among previously mentioned mechanisms, DNA methylation, is the most stable type of epigenetic modification that plays a significant role in regulating gene-expression, maintaining genomic integrity and influencing developmental processes and disease, in a cell-type specific manner (Armstrong et al 2019; Schubeler et al 2015; Smith et al. 2013). Specifically, it consists in the addition of a methyl group to the fifth carbon position of cytosine (5mC) (Kumar et al 2018; Suzuki eta al 2018). It is worthy to mention that DNA methylation exerts diverse regulatory effects depending on its genomic context, since it is not uniformly distributed across the genome. While hypermethylation of CpG rich promoters is linked to gene silencing in vertebrates (implicated in processes such as genomic imprinting and X-chromosome inactivation), global hypomethylation is associated with genomic instability and aberrant gene activation (contributing to disease development such as tumorigenesis, neurodegenerative diseases and neurological disorders (Hao et al 2017; Ou et al 2017; Younesian et al. 2024).

With the fast development of next-generation sequencing technologies, currently the most reliable and widely used method for measuring DNA methylation is bisulfite sequencing (BS-Seq), as it enables a detailed exploration of the epigenome at single-nucleotide resolution (Lister et al., 2009; Krueger et al. 2012). In particular, whole-genome bisulfite sequencing (WGBS) provides high-resolution map of DNA methylation across the entire genome, allowing for the identification of different methylation patterns genome-wide (Wulfridge P et al 2019). In this technique, DNA is treated with sodium bisulfite, converting unmethylated cytosine residues to uracil (and ultimately to thymine when PCR amplification occurs), whilst maintaining methylated cytosines unchanged and finally providing a single-base, quantitative cytosine methylation levels (Stirzaker C et al. 2014). However, the analysis of methylation sequencing data presents several computational challenges, including the increasing diversity of methodologies and library preparation protocols for WGBS, the complexity of bisulfite-converted data and the need for precise identification of differential methylated cytosines (DMCs) and differential methylated regions (DMRs) (Bock et al., 2011; Smallwood et al., 2011; Sun et al 2014).

The first step of analysing bisulfite-sequencing data is to align reads to a reference genome and to count the number of C-to-T conversions, using read mapping and methylation calling aligner tools, such us, *Bismark* (Krueger et al. 2011), *Methylcoder* (Pedersen et al. 2011), *BRAT* (Harris eta al 2010), *RMAP* (Smith et al 2009), *BSSeeker* (Chen et al 2010, Huang et al 2018) and *BSMAP* (Xi et al 2009), among others. Additionally, within the past few years, multiple computational approaches have been developed for downstream DNA methylation studies, which often involve detecting DMCs and DMRs (Wreczycka et al. 2017). Some of them implement linear regression designs, including *methylKit* (Akalin et al 2012), *RnBeads* (Assenov et al 2012) and *BSmooth* (Hansen et al 2012). Other commonly used software, are based in binomial model, for example, *edgeR* (Chen et al 2018) *DSS* (Feng et al 2014), *BiSeq* (Hebestreit et al 2013), *MOABS* (Sun et al 2014), *RADMeth* (Dolzhenko et al 2014), *Bisulfighter* (Saito et al 2014), *methylSig* (Park et al 2014), *DMRfinder* (Gaspar et al. 2017), *HMM-DM* (Yu et al. 2016).

Despite the improvements in tool development, a major challenge in DNA methylation analysis arises from the lack of a consensus on statistical and computational tools used for DNA methylation analysis. Indeed, there is no standard protocol to carry out this process (Liu et al 2020; Yu et al. 2016; Klein et al 2015). Specifically, given that these methods rely on distinct statistical models they often yield varying results, identifying different DMCs or different sets of DMRs with no large percentage of agreement. This variability complicates the interpretation of findings and raises concerns about the reproducibility and biological significance of the detected regions. In light of this inconsistency, and in order to have a deeper understanding of DMCs identification, in this article, we present and discuss the integration of two widely used tools to test predefined genomic regions for differential methylation in BS-seq data. On one hand, *methylKit* (Akalin et al 2012) and on the other one, *edgeR* (Robinson et al. 2010; Chen et al 2018), both are available within the statistical packages provided by the Bioconductor software on the R platform and have shown high sensitivity and/or low false positive rates. *methylKit* (Akalin et al 2012) applies a Fisher’s exact test or logistic regression with overdisperssion correction to calculate p-values adjusted to q-values for multiple test correction using *SLIM* approach (Wang et al 2011). While *edgeR* (Robinson et al. 2010; Chen et al 2018) is based on the linear model framework (*GLM)* (McCarthy et al. 2012) and models the variation between biological replicates through the negative binomial dispersion.

Our aim is to share a new perspective on the use of appropriate tool or tool combination for the identification of DMCs and differentially methylated genes (DMGs), while considering both biological and technical factors that influence methylation data analysis. In fact, it is expected that the combination of different advanced analytical tools will expand the frontiers of methylome analysis, with increasingly relevant applications in environmental epigenetic, biomedical research and precision medicine (Sun et al., 2021).

## METHODOLOGY

### Sample processing and DNA isolation procedures

Genomic DNA from control and morphine treated (10 μM, Alcaliber) mouse embryonic stem cells (mESCs; Oct4-GFP cell line / PCEMM08, PrimCells) was isolated and extracted, using a classic phenol-chloroform/isoamyl methodology with phenol (P4557, Sigma), chloroform (CL01981000, Scharlau) and isoamyl alcohol (BP1150, Fisher BioReagents) and following the manufacturer’s instructions. DNA concentration and purity were measured using the Nanodrop Spectrophotometer ND-1000 (Thermo Fisher Scientific) by quantifying the 260/280 absorbance ratio. The amount of extracted DNA from two samples was preserved at −80° prior to library preparation.

### Bisulfite conversion, library preparation and sequencing

Genomic DNA was sonicated using a Soniprep 150 to produce fragments of approximately 300 base pairs (bp). These fragments were subsequently denatured and prepared for bisulfite conversion, using the EZ DNA Methylation-lightning Kit to facilitate downstream methylation analysis. DNA fragments were then subjected to library preparation, employing the KAPA Library Preparation Kit along with xGenTM Methyl UDI-UMI Adapters. The quality and integrity of the libraries were assessed using an Agilent 2100 Bioanalyzer with the DNA 7500 assay. Following library preparation, sequencing was performed on an Illumina NovaSeq 6000 S4 platform (paired-end), with 4× multiplexing to ensure robust coverage, achieving a minimum of 50 000 reads per replicate for each sample. Reproducibility between the two biological replicates was assessed via Spearman correlation, demonstrating high concordance. All cytosine residues across both duplicates for each sample were included in subsequent genome-wide methylation analyses. The computational workflow summarizing all the steps is included in **Figure. 1**.

**Figure 1.**
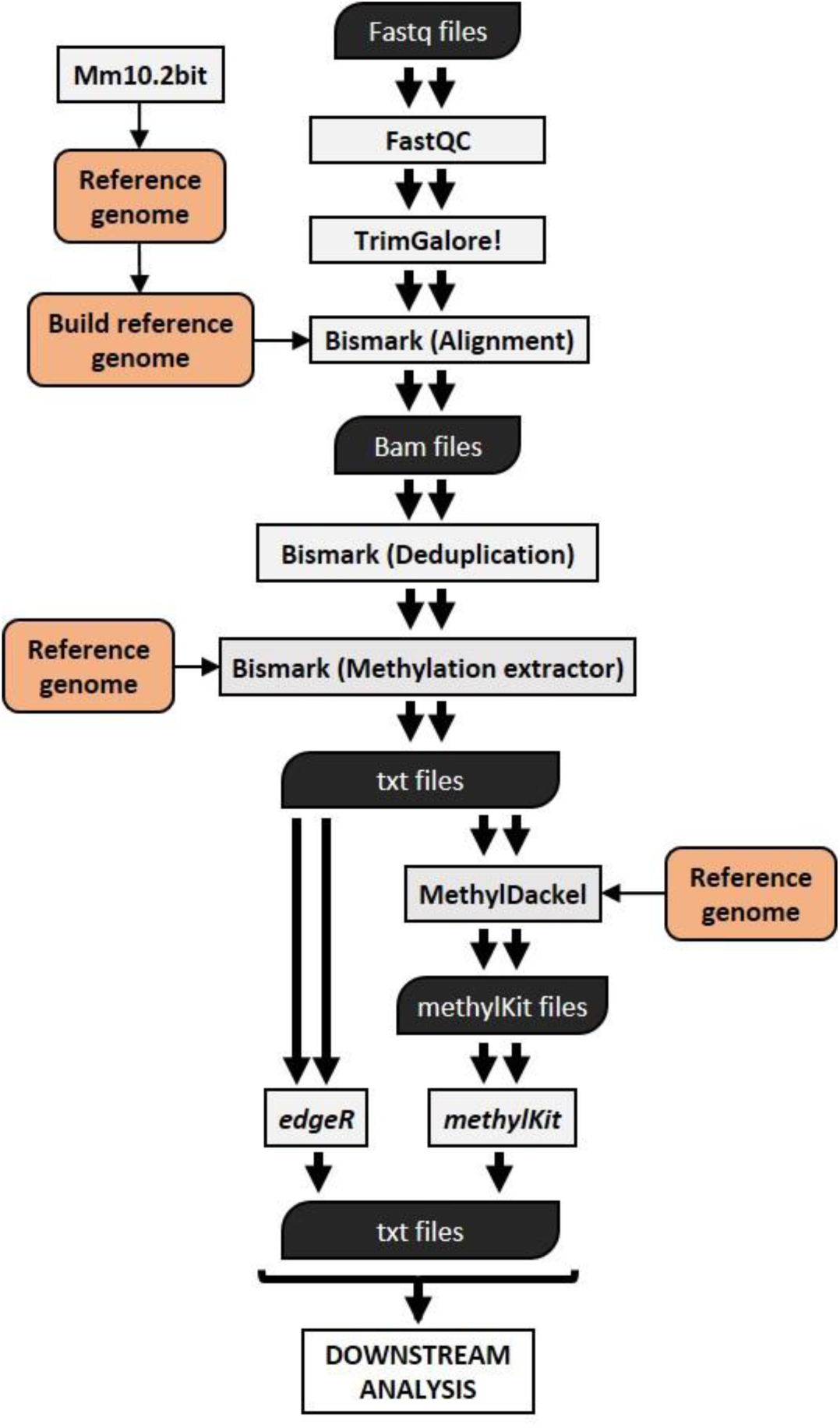
DNA methylation analysis workflow using bisulfite sequencing data. A summary of the followed protocol and file types is shown in a flow chart.

### Pre-processing

The library fragment adapters were trimmed using Trim Galore! (v.0.6.2) (Krueger F. 2019), and subsequently, a FastQC High-Throughput Sequence Quality Control report (v.0.11.6) (Andrews, 2010) was generated to evaluate the quality of the WGBS FASTQ files, indicating a quality score exceeding 30, and foreseeing a suitable mapping efficiency. The resulted reads were concatenated using *Cat* (v.8.22) (Torbjörn Granlund and Richard M. Stallman 2012). Spearman correlation confirmed the reproducibility of the two biological replicates in methylation samples (**Supplementary Figure 1A**).

### Cross platform analysis of m5C sequencing data

The WGBS data was mapped to bisulfite-converted reference genome file for mouse (*UCSC GRm38/mm1*0) with *Bowtie2* (Langmead and Salzberg 2012; Langmead eta al. 2019) on *Bismarck* (v.0.22.1) (Krueger and Andrews, 2011) with default parameters and the resulting binary alignment files were sorted and indexed with *SAMtools* (v.14.0) (Li et al. 2009) to enable quicker access. To be unable to use the post-processing programs that follow the WGBS bioinformatics analysis, *MethylDackel* (v.0.5.1) (MethylDackel 2020) tool was used to estimate the percentage coverage and percentage methylation for all CpG sites on a genome-wide scale. Then, to evaluate the effect of morphine on differentially methylated cytosines (DMC) two different statistical tools were chosen: 1) *edgeR* (v.3.32.1) (Robinson et al 2010; Chen et al 2018), which uses the negative binomial distribution to assess statistical significance, with TMM normalization accounting for library size. 2) *methylKit* (v.1.16.1) (Akalin et al 2012), which normalizes read coverage distribution between samples. In both cases regions with a p-value < 0.05 were considered differentially methylated. The PCA (*principal component analysis*) in each of the tools confirmed the similarity between replicates and distinguishable differences between samples (control and morphine treated duplicate samples). After applying TMM normalization (correcting for library size), read count distribution plot showed a correct data normalization and *Volcano plot* analysis confirmed the differences in these DMCs analyzed through two tools (**Supplementary Figure 1B**). It is worthy to mention that both tools present the DMCs in different ways: *edgeR* uses fold change values and *methylKit* applies percentage values. The methylation landscape is viewable via *UCSC genome browser* (Kent et al 2002). Integrative analyses between both tools were performed using *Venny tools* (v.2.1.0) (Oliveros JC et al 2007-2015), and *The Gene Ontology Resource from the GO Consortium* (https://geneontology.org/, v.16.1.0) (Ashburner et al. 2000; The Gene Ontology Consortium 2019) was used to identify the biological functions.

## RESULTS

### Overview of DMR detection tools/ similarity analysis of DMR detection tools

First of all, a methodological comparative analysis was performed between both R packages *edgeR* and *methylKit*. **Supplementary Table 1** summarizes the main features of each tool used for analysing DMCs (including platform, source and implementation language, alignment ability, single or paired end capacity, major function, concept, model or test, statistical method, quality control and preprocessing step, smoothing and further analysis options). Both packages are compatible with Windows and Linux platforms, and although neither includes alignment functionality, both support input from single-end and paired-end sequencing data, making them versatile in terms of sequencing protocols. Importantly, both tools support differential methylation analysis, even if using different statistical and conceptual frameworks.

Differential DNA methylation is usually calculated by comparing the proportion of methylated cytosines in a test sample relative to a control. In terms of used concept and model or tests, *edgeR* is primarily designed for RNA-seq but can be adapted for DNA methylation data when formatted as count data. It utilizes a negative binomial regression model that accounts for overdispersed data, and supports both competitive and self-contained tests. Statistical inference in *edgeR* can be carried out using generalized linear model likelihood ratio tests (GLM LRT), quasi-likelihood F-tests (QLF), or an exact test that is conceptually similar to Fisher’s exact test, but tailored for overdispersion. In contrast, *methylKit* is specifically designed for bisulfite sequencing data and the methylation levels of all samples at a given CG site are modelled by a logistic regression. It applies a self-contained testing approach, using either Fisher’s exact test or logistic regression to compute p-values for DMCs.

Systematic and base-calling errors can impact DMCs identification, making quality control of raw methylation data essential, particularly in coverage terms. In this sense, both tools implement quality control and preprocessing strategies to mitigate technical biases*. edgeR* performs coverage normalization and removes low-coverage regions to ensure accurate modelling of count data. Similarly, *methylKit* applies library size normalization and uses similar criteria for filtering and removing potential duplication bias, enhancing the reliability of downstream statistical testing. Furthermore, due to the similarity of methylation levels among neighbouring CG sites, smoothing algorithms help estimate values for low-coverage or uncovered sites and reduce sequencing errors, though they may introduce some bias. Notably, neither tool incorporates smoothing techniques for methylation data, although both support further downstream analysis to add genomic context to the obtained results. For example, both tools allow hyper- and hypo-methylation global levels and events per chromosome, web-based methylation data visualization (UCSC Genome Browser and Integrated Genome Viewer) and genetic annotation, aiding in biological interpretation of DMCs.

### Similarity analysis of DMC detection tools

To evaluate the concordance between *edgeR* and *methylKit* in the detection of DMCs, both tools were applied to DNA methylation data derived from control and morphine treated mESCs samples. *edgeR* identified a total of 203337 DMCs, while *methylKit* detected 223280 DMCs. To further characterize these findings more in depth, changing trends in DMCs were identified. Of these, *edgeR* classified 60704 DMCs as hypermethylated (29.85%) and 142633 as hypomethylated (70.15%). Similarly, *methylKit* identified 67054 hypermethylated DMCs (30.03%) and 156226 hypomethylated DMCs (69.97%) (**Table 1 and Figure 2A**). Then the overlap between DMCs detected by both tools was identified, rising to the amount of 153 394 DMCs, corresponding to 56.14% of the total number of DMCs. Within this overlapping set, 44 641 common DMCs (29.10%) were classified as hypermethylated, while 108 753 common DMCs (70.90%) were identified as hypomethylated. This was equivalent to a 53.71% and a 57.2% fraction of overlapped DMCs respectively (**Table 1 and Figure 2A**). Additionally, the mean and the standard deviation were calculated for both tools, to further estimate the coefficient of variation. This value was less than 5% (0.05) in all cases, indicating low relative variability. In order to focus in the common identifications, the distribution of overlapping DMCs was analysed for each chromosome. To explore the chromosomal distribution of commonly identified DMCs, their frequency across chromosomes was analysed. The majority of the DMCs were significantly located on chromosomes 8 and 11. In contrast, chromosomes 2, 3, 4, 7, 12, 13, 18, 19 and sex chromosomes (X and Y) exhibited a notably lower number of DMCs. However, it is worthy to mention that a consistent pattern was observed across all chromosomes, the number of hypomethylated cytosines was always higher than hypermethylated ones (**Supplementary Figure 2**).

**Figure 2.**
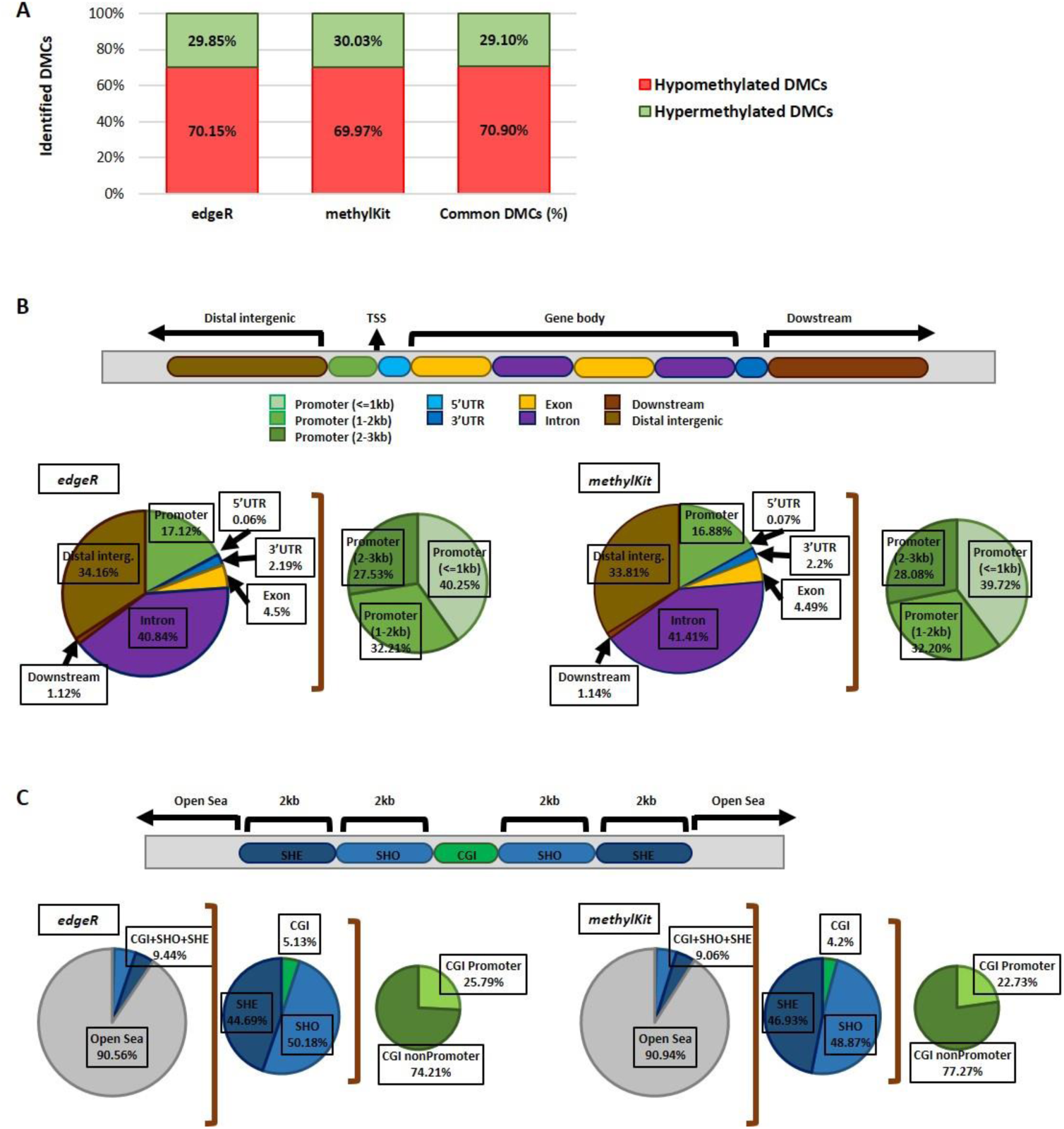
Distribution of DMCs across different genomic regions. (A) Column chart displaying the percentage of identified hypermethylated and hypomethylated DMCs for each tool and commonly identified overlapped sites. (B) Piechart showing DMCs distribution around gene features, in promoter (divided in <=1kb, 1-2 kb and 2-3 kb), 5’UTR, 3’UTR, exon (1^st^ and others), intron (1^st^ and others), downstream of the gene end and intergenic regions, for *edgeR* (left) and *methylKit* (right). (C) Pie chart showing DMCs distribution around different CpG-related features, in CpG island (belonging to promoter or nonpromoter region+1 kb from TSS), Shore (<2kb), Shelf (<4kb) and Open sea (rest of the genome) for *edgeR* (left) and *methylKit* (right).

**Table 1.** Comparison of identified DMCs between *edgeR* and *methylKit*. Total, hypermethylated and hypomethylated DMCs are specified for *edgeR* and *methylKit*, together with common sites identified by both tools.

| | *edgeR* | *methylKit* | $\bar{x}$ | $\sigma$ | $C_v$ (%) | Common DMCs |
| --- | --- | --- | --- | --- | --- | --- |
| Identified total DMCs | 203 337 | 223 280 | 213308.5 | 9971.5 | 4.67 | 153 394 (56.14%) |
| Hypermethylated DMCs | 60 704 | 67 054 | 63879 | 3175 | 4.97 | 44 641 (53.71%) |
| Hypomethylated DMCs | 142 633 | 156 226 | 149429.5 | 6796.5 | 4.55 | 108 753 (57.21%) |

Given the functional relevance that DNA methylation could have on gene expression, the next step was to examine each tool’s capacity in detecting DMCs across specific gene-associated features. Particular attention was given to DMCs located in promoter regions, due to their central role in transcriptional regulation. The following gene features were included in the analysis: promoters (<=1 kb, 1-2 kb and 2-3 kb upstream of transcription start sites), 5’UTR, 3’UTR, exons, introns, downstream regions and distal intergenic regions (**Figure 2B**). In promoter region *edgeR* identified a total of 34817 DMCs (14015 DMCs within <=1 kb, 11216 DMCs in 1-2 kb and 9586 DMCs in 2-3 kb), while *methylKit* detected a total of 37699 DMCs (14974 DMCs within <=1 kb, 12140 DMCs in 1-2 kb and 10585 DMCs in 2-3 kb). Identified number of 5’UTR and 3’UTR DMCs were very similar in both tools, specifically 129 and 4460 DMCs in *edgeR* and 149 and 4920 in *methylKit*. Furthermore, for exonic and intronic regions, *edgeR* reported 9146 and 83043 DMCs, whereas *methylKit* identified 10024 and 92460 DMCs respectively. Similarly, for downstream and distal intergenic regions, *edgeR* identified 2284 and 69458 DMCs, while *methylKit* reported 2540 and 75488 DMCs respectively. As shown in **Figure 2B**, most DMCs were located in introns and distal intergenic regions, corresponding to 40.84% and 34.16% of gene features in *edgeR*, and conforming 41.41% and 33.81% in *methylKit*. Promoter regions and adjacent areas collectively accounted for approximately 17% of DMCs in both tools. In any case, the distribution of DMCs across genomic features was consistent between tools, with only minor variations (**Table 2 and Figure 2B**). Then, to asses overlap between both tools, common DMCs within specific gene features were identified. For promoter region a total of 26031 DMCs were identified (10324 DMCs within <=1 kb, 8412 DMCs in 1-2 kb and 7295 DMCs in 2-3 kb); 100 DMCs in 5’UTR; 3403 DMCs in 3’UTR; 6909 DMCs in exons; 63657DMCs in introns; 1699 DMCs in downstream regions; and 51595 DMCs in distal intergenic regions. In all cases, the fraction of overlapped DMCs remained close to 56%, in the same way as observed in the general dataset. Additionally, the coefficient of variation remained below 5% (0.05) in the majority of cases, reinforcing the reproducibility of the findings (**Table 2**).

**Table 2.**
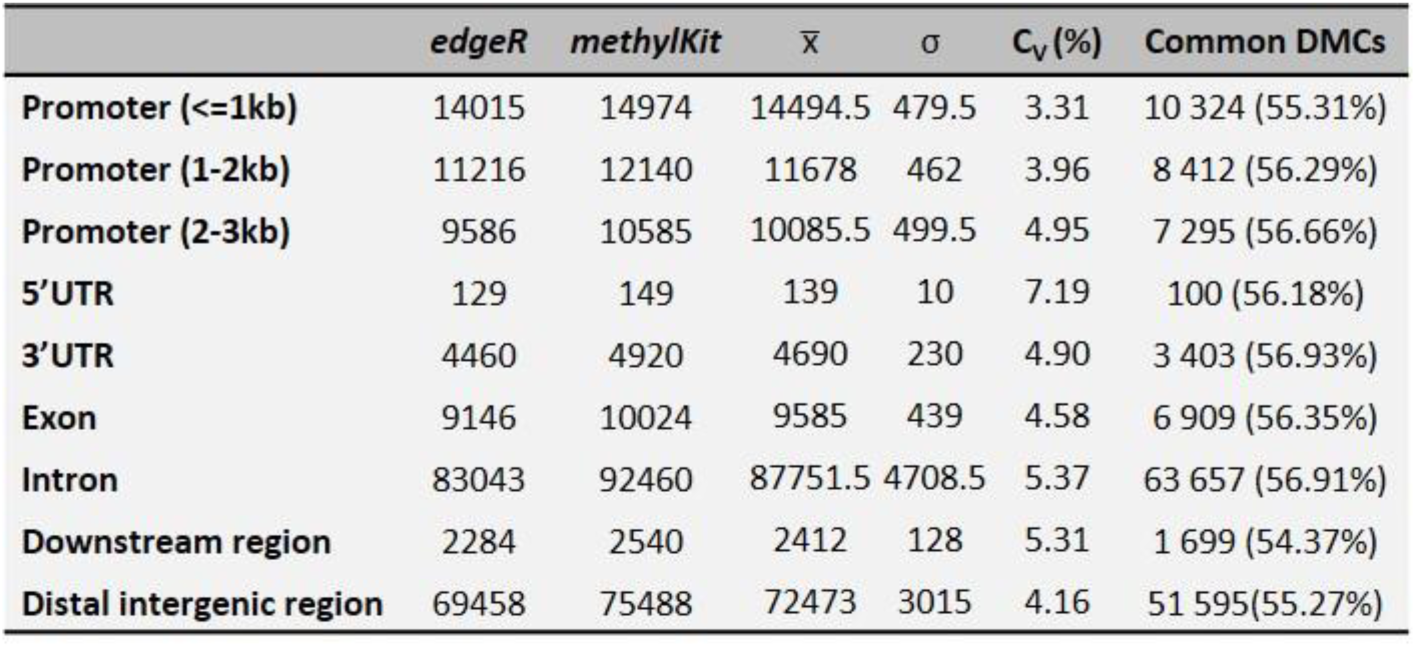
Comparison of identified gene features between *edgeR* and *methylKit*. DMCs belonging to promoter (divided in <=1kb, 1-2 kb and 2-3 kb), 5’UTR, 3’UTR, exon (1^st^ and others), intron (1^st^ and others), downstream of the gene end and intergenic regions are specified for *edgeR* and *methylKit*, together with common sites identified by both tools.

DNA methylation is not uniformly distributed across the genome. Instead, CpG sites tend to cluster in distinct genomic contexts, including CpG islands (CGIs), shores, shelves and open sea regions (**Figure 2C**), each with potentially different genetic regulatory roles. Consequently, the way forward was to evaluate the ability of each tool to identify DMCs within these CpG-related features. Both tools predominantly identified DMCs in the area known as open sea (over 90% of all DMCs), that is, *edgeR* detected 184129 DMCs and *methylKit* identified 203048 DMCs outside the CpG islands area. Likewise, *edgeR* identified 985 DMCs in CGI, 9638 DMCs in shores and 8585 DMCs in shelves (summing up to 9.44%), while *methylKit* detected 849 DMCs in CGI, 9888 DMCs in shores and 9495 DMCs in shelves (amounting to 9.06%). In fact, from the 1% of DMCs located in CGIs that simultaneously coincided with promoter regions ranged from 25.79% in *edgeR* to 22.73% in *methylKit* (**Table 3 and Figure 2C**). Next, overlap analysis revealed 566 common DMCs in CGIs, 6839 DMCs in shores, 6604 DMCs in shelves and 139385 in open sea regions. As with previous comparisons, the proportion of overlapping DMCs remained close to 56% across all categories, except for CGIs, which exhibited a lower overlap of 44.64%. A coefficient of variation under 5% (0.05) was observed in the vast majority of cases, suggesting again low variability between replicates (**Table 3**).

**Table 3.**
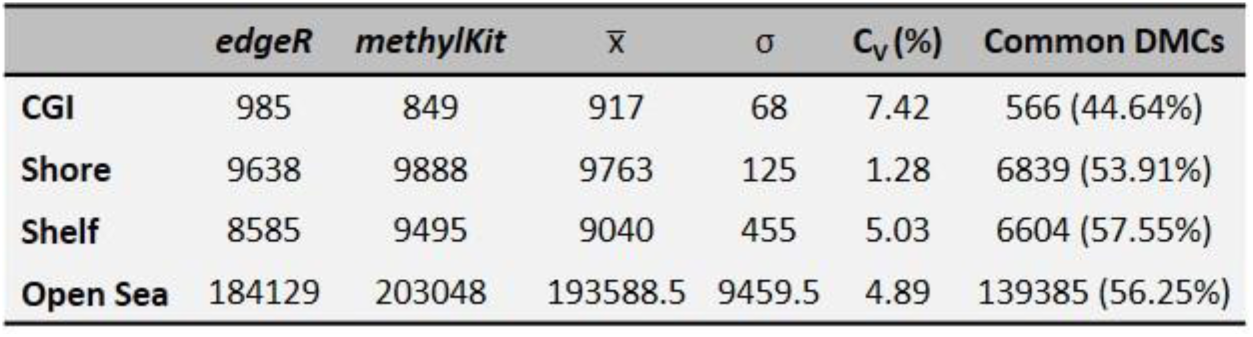
Comparison of identified CpG-related features between *edgeR* and *methylKit*. DMCs belonging to CGI, Shore (<2kb), Shelf (<4kb) and Open sea (rest of the genome) regions are specified for *edgeR* and *methylKit*, together with common sites identified by both tools.

### Similarity analysis of DMG detection tools

When identifying DMCs, it is crucial for a tool that facilitates the identification of DMGs, as these provide a more biologically meaningful and functionally relevant perspective. While DMCs offer site-specific information on methylation changes, DMGs enable the contextualization of these modifications at the gene level, linking methylation patterns to biological processes, transcriptional effect and thereby facilitating interpretation within physiological and pathological context. Therefore, a comparative evaluation of both tools in DMG detection was conducted. *edgeR* identified a total of 17657 DMGs, whereas *methylKit* reported 17772 DMCs. Within these, *edgeR* classified 13128 DMGs as hypermethylated, 16313 DMGs as hypomethylated and 11783 DMGs with both alterations (66.7%). Similarly, *methylKit* identified 13426 hypermethylated DMGs, 16429 hypomethylated DMGs and 12081 with both marks (68%) (**Table 4 and Figure 3A**). Analysis of the overlap revealed 16357 DMGs detected by both tools, corresponding to 87.8% of all DMGs. Of these, 11594 were commonly classified as hypermethylated DMGs, and 14954 as hypomethylated DMGs, while 10191 common DMGs displayed both methylation alterations (62.3%). This was equivalent to 80%, 86.2% and 74.53% of the overlapping DMGs respectively (**Table 4 and Figure 3A**). In all comparisons, the coefficient of variation remained below 5% (0.05), reinforcing the reproducibility of the observations (**Table 4**).

**Figure 3.**
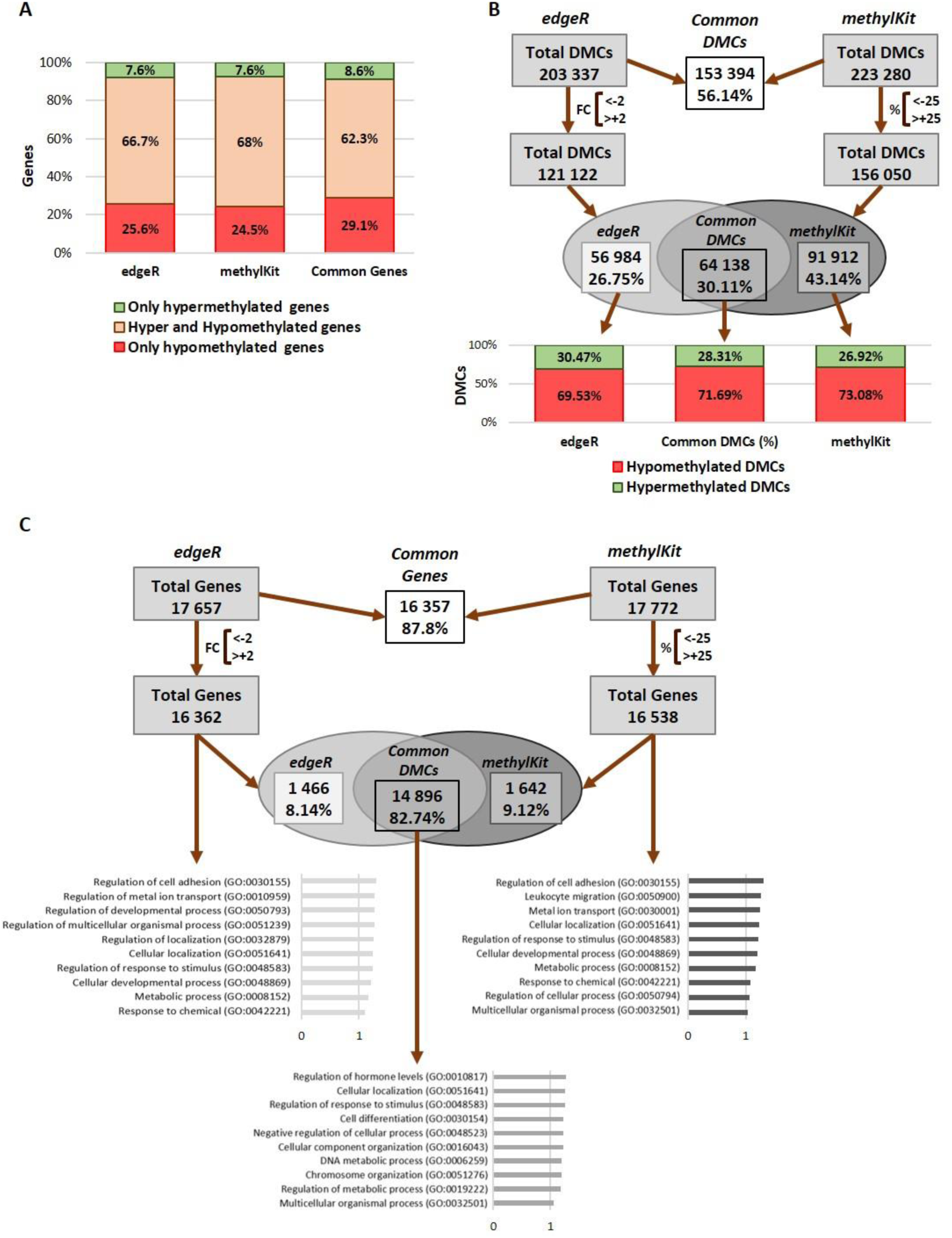
Identification of biologically relevant results. (A) Column chart displaying the percentage of only hypermetylated, only hypomethylated and hyper/hypomethylated genes for each tool and commonly identified overlapped sites. (B) Total number of DMCs identified before and after the applied threshold for each tool (x<-2 and x>+2 fold change for *edgeR* and x<-25 and x> +25 percentage for *methylKit*), Venn diagram showing the overlap between common DMCs sites and a percentage column chart displaying hyper and hypomethylated DMCs proportions. (C) Total number of differentially methylated genes identified before and after the applied threshold for each tool (x<-2 and x>+2 fold change for *edgeR* and x<-25 and x> +25 percentage for *methylKit*), Venn diagram showing the overlap between common genes and a functional enrichment analysis showing the most indicative biological functions of the specific genes annotated from each tool. Statistical analyses Bonferroni corrected for p<0.05.

**Table 4.** Comparison of identified differentially methylated genes between *edgeR* and *methylKit*. Total, hypermethylated, hypomethylated and hyper/hypomethylated genes are specified for *edgeR* and *methylKit*, together with common sites identified by both tools.

| | *edgeR* | *methyKit* | $\bar{x}$ | $\sigma$ | $C_v(\%)$ | Common Genes |
| --- | --- | --- | --- | --- | --- | --- |
| Identified total Genes | 17 657 | 17 772 | 17 714.5 | 57.5 | 0.32 | 16 357 (87.8%) |
| Hypermethylated Genes | 13 128 | 13 426 | 13 277 | 149 | 1.12 | 11 594 (80%) |
| Hypomethylated Genes | 16 313 | 16 429 | 16 371 | 58 | 0.35 | 14 954 (86.2%) |
| Hyper and Hypomethylated Genes | 11 783 | 12081 | 11932 | 149 | 1.25 | 10191 (74.53%) |

Significantly, while the general overlap in DMCs between *edgeR* and *methylkit* was 56%, this value increased substantially to 87.8% when focusing on the gene level (DMGs). To enhance the biological relevance of the results and minimize false positives, equivalent stringency thresholds were applied. That is, a minimum fold-change difference of 2 in *edgeR* and minimum methylation difference of 25% in *methylKit* (with significance set at p ≤ 0.05 in both cases) (**Figure 3B and C**). These parameters reduced the total number of DMCs to 121122 in *edgeR* and 156050 for *methylKit*. The resulting intersection of both tools identified 64138 common DMCs, corresponding to 30.11% of the total. Despite this reduction, the distribution of methylation trends remained consistent with previously described results, since common hypermethylated DMCs were 28.31% and common hypomethylated DMCs were 71.69% (**Figure 3B**). Focusing on the genetic impact, these thresholds decreased the number of DMGs to 16362 for *edgeR* and 16538 for *methylKit*, with an overlap of 14896 DMGs. It is worthy to mention that this overlap was 82.74% of the total number of DMGs. To measure the biological relevance of the identified DMGs, Gene Ontology enrichment analyses were performed separately on the DMGs identified by each tool, as well as on the set of DMGs commonly detected by both tools, revealing substantial biological overlap. Both *edgeR* and *methylKit* identified processes related to cellular development, metabolic activity, chemical stimulus response, cellular localization and regulation of cell adhesion, reflecting core biological functions across datasets. The common list reinforced these findings by highlighting biological processes such as cellular localization, regulation of metabolic and cellular processes, regulation of response to stimulus, and multicellular organismal processes, thereby emphasizing the robustness and the biological relevance of the shared results and offering complementary insights suitable for integrative interpretation (**Figure 3C**).

## DISCUSSION

This study provides a comprehensive evaluation and comparison analysis between *methylKit* and *edgeR*, two widely used computational tools for detecting DMCs in WGBS data. In brief, both methods proved effective in identifying DMCs and different genomic contexts, showing that both tools can be complementary; although notable differences were observed in the total number detected and the way the results were presented. These findings highlight the importance of considering methodological integration in methylation analyses and underscore the need for standardized protocols to ensure reproducibility and consistency of results in biological relevance.

Although *methylKit* and *edgeR* share similar objectives, they differ considerably in their statistical frameworks and normalization strategies. *edgeR* employs a generalized linear model based on a negative binomial distribution with empirical Bayes dispersion estimation and TMM (Trimmed Mean of M-values) normalization (which may reduce false positives) (Chen et al. 2018), while *methylKit* implements Fisher’s exact test or logistic regression with overdispersion correction, applying multiple testing correction via the SLIM method (Akalin et al. 2012). These methodological differences were reflected in how each tool reported DMCs, as *methylKit* provided percentage change values, whereas *edgeR* presented fold change values. However, this did not present any significant limitation when comparing the obtained data, nor in the establishment of an equivalent threshold between both tools to identify the most relevant methylation changes (Akalin et al. 2012; Chen et al. 2018). In contrast, it is worthy to mention that, *methylKit* identified more total DMCs than *edgeR* (223280 vs 203337 DMCs), which on one hand could be potentially related to the correction for overdispersion in normalization that may lead to a more conservative DMC detection in *methylKit*. And on the other hand, that identification difference may reflect *methylKit* design as a methylation-specific tool (Akalin et al. 2012), unlike *edgeR,* which was initially designed to analyse RNA-seq (Robinson et al. 2010) and later modified to identify methylation changes (Chen et al. 2018). These differences support previous findings indicating that the choice of analytical pipeline can significantly affect both the sensitivity and specificity of methylation calls (Ziller et al. 2015; Liu et al. 2020). Importantly, the cross-platform comparison revealed a moderate level of concordance (56.14%) in DMC detection, with a coefficient of variation below 5%, indicating a strong technical robustness and reproducibility of results, even if being a dataset with a limited number of replicates. This result agreed with earlier benchmarking studies comparing methylation tools, which have noted similarly limited overlap among DMC detection methods using distinct statistical models (Liu et al. 2020; Ziller et al. 2015). Despite these differences, both tools exhibited a shared methylation directionality trend, with a predominant hypomethylation pattern (approximately 70% of DMCs), not only globally but also across individual chromosomes. Therefore, these results suggest that tool integration remains critical to identify robust biological signals in methylome studies.

The distribution of DMCs was largely consistent between *edgeR* and *methylKit*, particularly within intronic and distal intergenic regions, which together represented over 70% of total DMCs. This distribution agrees with previous epigenomic landscape analyses showing that DNA methylation variability occurs widely distributed throughout the genome (not confined to promoter regions) and may regulate enhancer activity and long-range chromatin interactions (Roadmap Epigenomics Consortium 2015; Kulis & Esteller et al. 2010; Maunakea et al. 2013; Schübeler et al. 2015). When looking to the rest of the gene features, both tools demonstrated similar distribution, further underscoring their robustness despite differences in statistical methodology. Interestingly, although promoter regions accounted for a smaller proportion of total DMCs in both tools (∼17%), this result has mechanistic relevance, since promoter methylation is linked to gene silencing (Bird, 2002; Deaton and Bird, 2011). This trend also was followed by DMCs distributed across different CpG context. Specifically, a minor fraction mapped to CGIs, shores and shelves in both tools and the vast majority of identified DMCs resided in open sea regions (over 90%). These results reinforce the evidence that open sea methylation may act as a dynamic regulatory element sensitive to epigenetic modulation (Hansen et al., 2011). Although DMCs within CGIs were less frequent, they displayed a high co-occurrence with promoter regions (approximately 25% of CGI DMCs overlapped with promoters), highlighting their potential regulatory relevance. This is in line with the view that methylation outside of canonical CGIs, particularly in enhancer regions, may provide more sensitive markers of cell state and environmental perturbation (Greenberg & Bourc’his, 2019). These results suggest that while both tools can detect DMCs in high-density CpG contexts, their primary sensitivity lies in identifying methylation variation in lower-density regions. These results align with studies suggesting that integrative approaches, combining multiple algorithms, can provide the most reliable view of methylation landscapes (Liu et al 2020; Yu et al. 2016; Klein et al 2015).

In terms of biological relevance, the most highlighting evidence for complementarity between the tools was identified from the gene level analysis. Despite the moderate overlap in DMCs, both *edgeR* and *methylKit* identified a high concordance (87.8%) in DMGs identification. This indicates that the biological impact of methylation changes is preserved at the gene level, even if the precise cytosine positions differ between tools. Furthermore, gene ontology enrichment analyses of the DMGs, underlined processes related to stimulus response, cell adhesion, metabolism and development pathways, in each tool separately, but also after overlapping results, functions that are known to be affected by opioid exposure in other studies (Hasin et al. 2017; Browne et al 2020; Walker et al. 2021). Importantly, the observed increase in overlap when analysing DMGs (compared to DMCs) suggests that gene-level aggregation may reduce false positives and increase biological interpretability, which supports its use as a complementary tool in methylation studies (Jaffe et al., 2012).

While this study provides valuable insights into the comparative performance of two key tools, several limitations should be acknowledged. First, the analysis was restricted to one experimental model (mESCs, single cell type and one treatment condition, morphine treatment), that is why results may differ in other biological contexts or with different exposure durations and doses. Second, we focused on DMCs and DMGs rather than DMRs, thereby, future work should consider DMRs, which may provide additional site-level information worthy to take into account. Third, the inclusion of only two replicates per condition may limit statistical power and increase sensitivity to outliers. Future studies should incorporate more replicates and explore integrative frameworks that combine WGBS with transcriptomic or proteomic data for functional validation.

Taken together, these findings underscore the benefits of integrative and complementary strategies to provide a more comprehensive picture of methylation dynamics. While no single tool can provide a complete and accurate representation of the methylome, integrating results from multiple statistical approaches can strengthen analytical confidence and enable more reliable identification of functionally relevant regions, thus reinforcing biologically relevant interpretation. Future directions should involve benchmarking these and other tools across a broader range of biological systems and experimental designs, to establish more standardized protocols and promote reproducibility in methylomic research.

## CONCLUSION

In conclusion, this comparative study demonstrates that while *edgeR* and *methylKit* each have unique advantages, their complementary use enhances the robustness and biological interpretability of differential DNA methylation analysis. The convergence in general trends, specifically the predominance of hypomethylation in response to morphine and the regional consistency of DMCs across gene features, supports the validity of findings derived from both statistical frameworks. Additionally, the high concordance of overlapped DMGs identification among both tools, highlights not only methodological consistency but also reinforces the biological relevance of the shared results.

Given the lack of a standardized pipeline in the field, cross-validation across multiple platforms remains essential to ensure improving accuracy in DMC detection and robust and reproducible epigenetic profiling. As DNA methylation continues to gain visibility as a key epigenetic biomarker, computational strategies that combine sensitivity, specificity, and flexibility will be essential. Future studies should continue exploring the impact of pipeline design, normalization strategy, and statistical modeling in large-scale WGBS datasets, to enable more comprehensive and biologically meaningful methylome interpretations.

## ACKNOWLEDGMENTS

The authors particularly acknowledge SGIKer resources of UPV/EHU for technical support with the computational calculations, which were carried out in the Arina informatics cluster.

## AUTHOR’S CONTRIBUTION

IMH, MA, IC and MA carried out the experiments and generated data. IMH performed the bioinformatic analysis. NS designed the study. NS and IMH discussed the data and IMH wrote the original manuscript. All the authors contributed to the writing of the manuscript, made critical comments and approved the final version.

## AVAILABILITY OF DATA AND MATERIAL

Sequencing data have been deposited into the Gene Expression Omnibus (GEO) under the accession number GEO: GSE292082 (https://www.ncbi.nlm.nih.gov/geo/query/acc.cgi?acc=GSE292082).

---

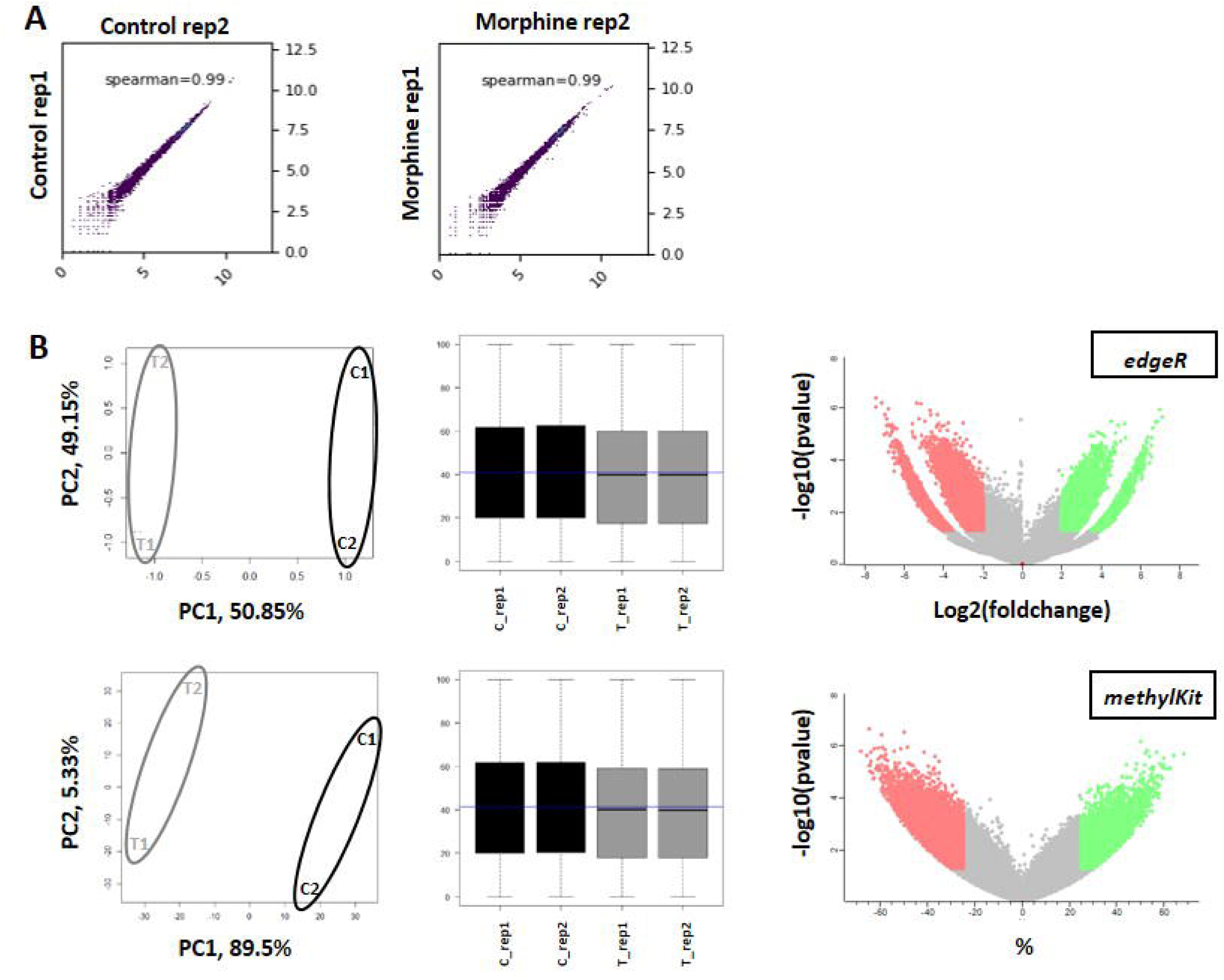

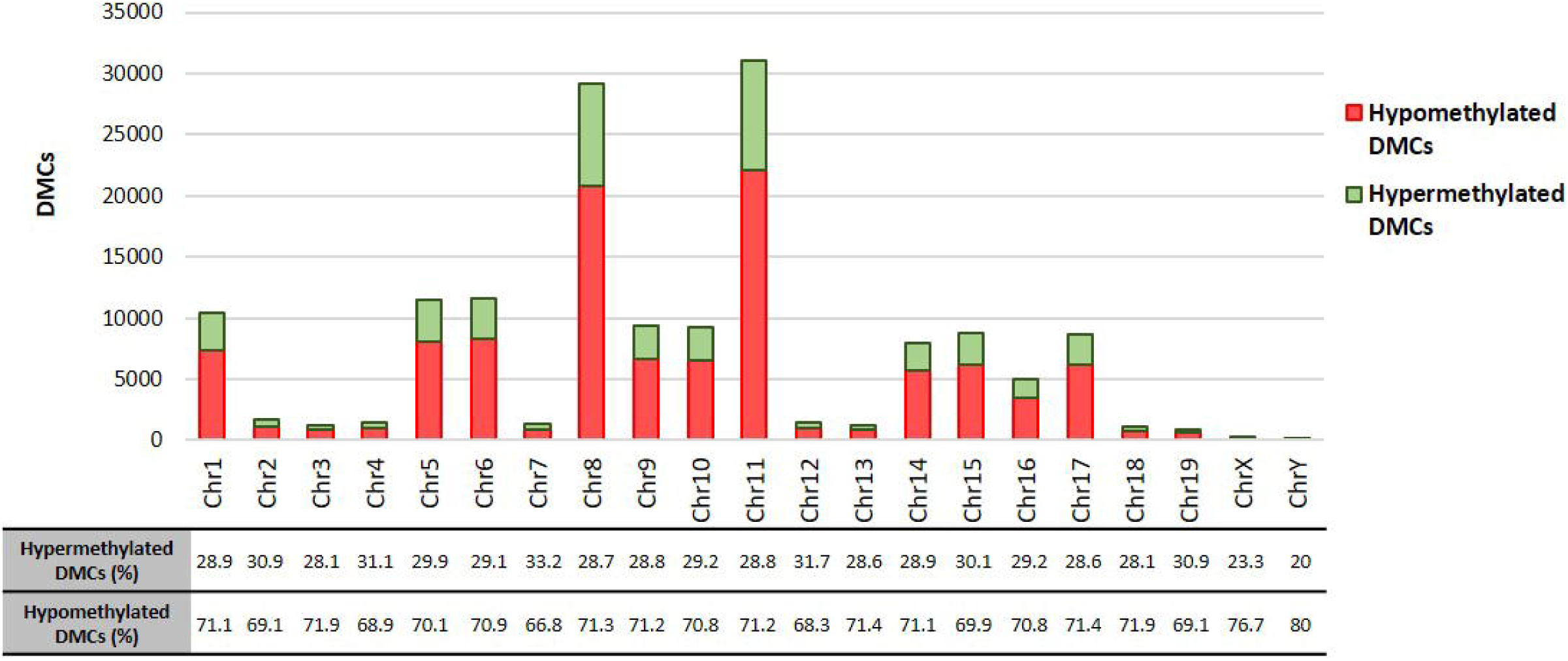

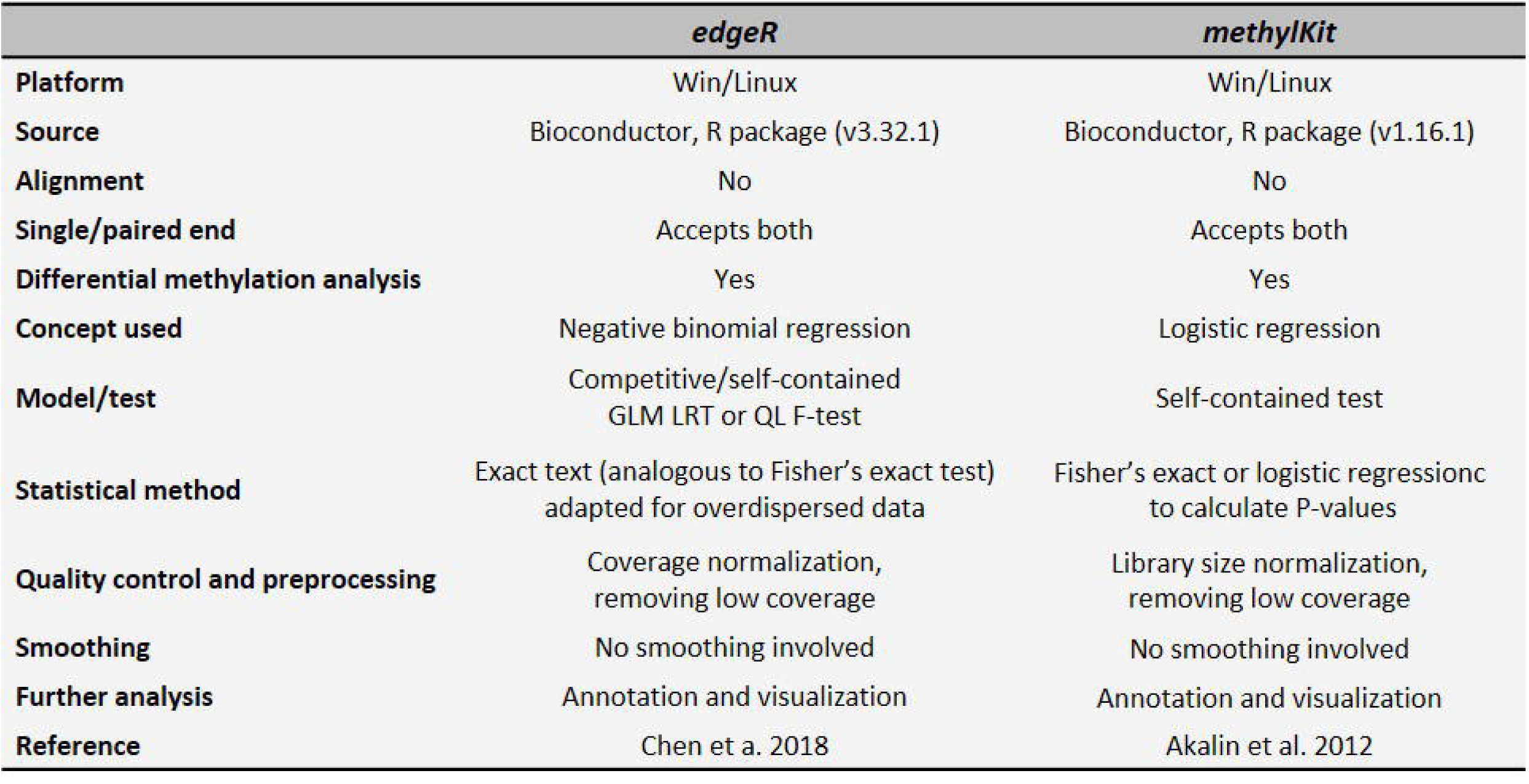

