## Supplementary material for "A Comparative Assessment of *edgeR* and *methylKit* Pipelines for DNA Methylation Detection"

**Affiliations:**
(1) Nursing I Department, Faculty of Medicine and Nursing, University of the Basque Country, Leioa, Bizkaia, Spain.
(2) Department of Physiology, Faculty of Medicine and Nursery, University of the Basque Country, Leioa, Bizkaia, Spain.
(3) Bizkaia Health Research Institute, Barakaldo, Bizkaia, Spain.

*To whom correspondence should be addressed:

Iraia Muñoa Hoyos, Nursing I Department. Faculty of Medicine and Nursing. University of Basque Country. 48940. Leioa, Bizkaia, Spain. +34 946015536.

**ORCID:**
Iraia Muñoa-Hoyos: 0000-0002-5107-6991
Manu Araolaza 0000-0003-1838-4594
Irune Calzado: 0000-0002-1787-6792
Nerea Subirán: 0000-0001-9202-1287

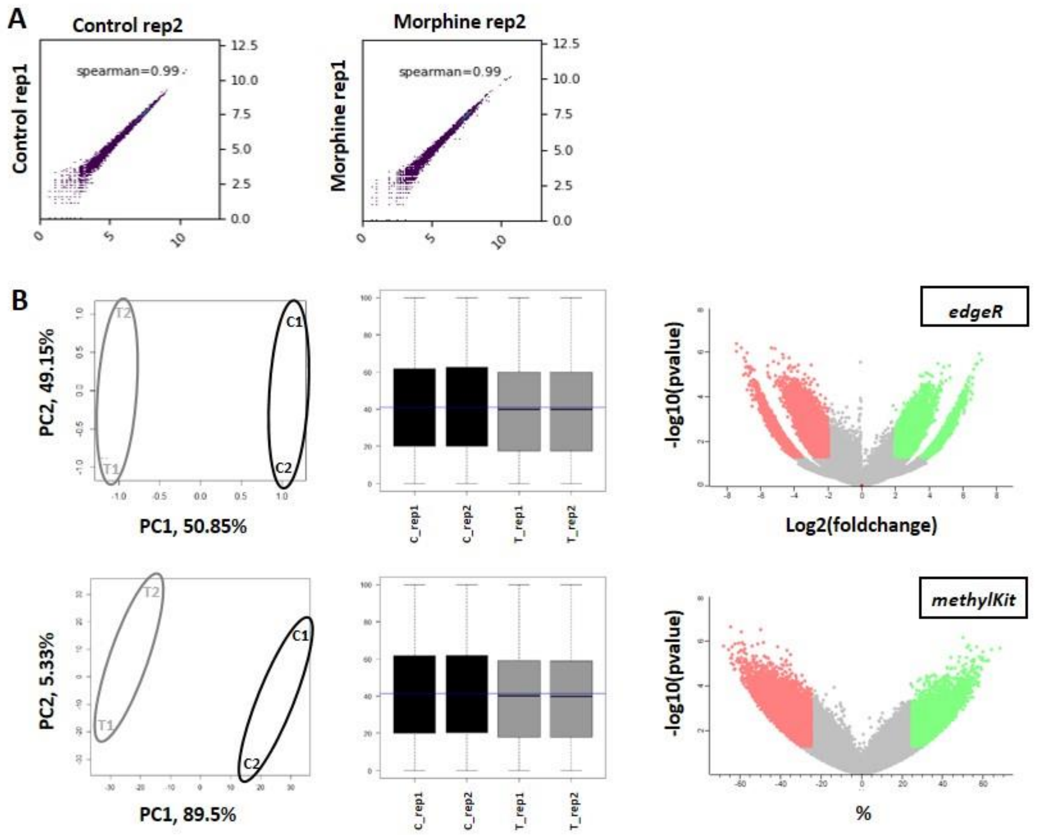


**Supplementary Figure 1. Summary of sample analysis.** (A) *Spearman* correlation analysis to analyze the reproducibility of sample types in Control samples (left) and Morphine treated samples (right). (B) *Principal Component Analysis* (PCA) *plot* to analyse similarities and differences between samples, normalization of data used in PCA analysis and *Volcano plot* analysis representing *Differentially Methylated Cytosines* (DMC) obtained by *edgeR* (superior) and *methylKit* (inferior). For the *Volcano plot,* the red dots indicate hypomethylated DMCs, and the green dots, hypermethylated DMCs ($p<0.05$). Gray dots were not significant ($p<0.05$), so they were rejected.

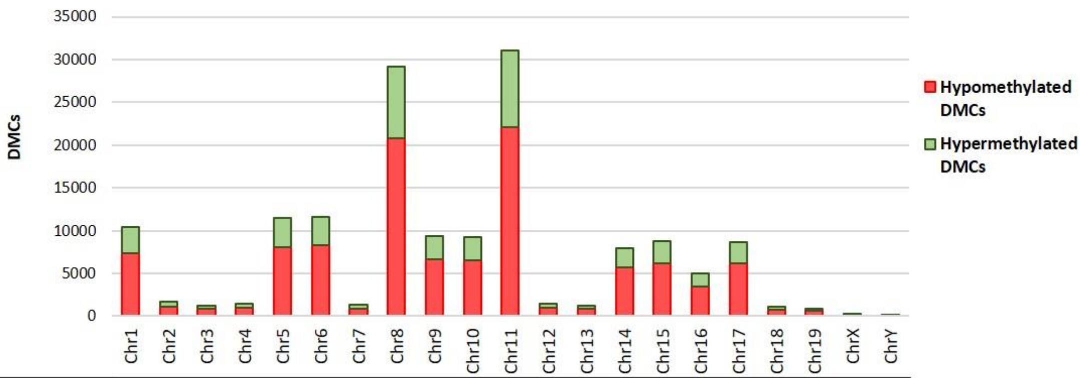


| | Chr1 | Chr2 | Chr3 | Chr4 | Chr5 | Chr6 | Chr7 | Chr8 | Chr9 | Chr10 | Chr11 | Chr12 | Chr13 | Chr14 | Chr15 | Chr16 | Chr17 | Chr18 | Chr19 | ChrX | ChrY |
|---|---|---|---|---|---|---|---|---|---|---|---|---|---|---|---|---|---|---|---|---|---|
| Hypermethylated DMCs (%) | 28.9 | 30.9 | 28.1 | 31.1 | 29.9 | 29.1 | 33.2 | 28.7 | 28.8 | 29.2 | 28.8 | 31.7 | 28.6 | 28.9 | 30.1 | 29.2 | 28.6 | 28.1 | 30.9 | 23.3 | 20 |
| Hypomethylated DMCs (%) | 71.1 | 69.1 | 71.9 | 68.9 | 70.1 | 70.9 | 66.8 | 71.3 | 71.2 | 70.8 | 71.2 | 68.3 | 71.4 | 71.1 | 69.9 | 70.8 | 71.4 | 71.9 | 69.1 | 76.7 | 80 |

**Supplementary figure 2. Genomic distribution of commonly identified DMCs.** Column chart displaying total number of DMCs in each cromosome commonly identified between *edgeR* and *methylKit* together with hypermethylated and hypomethylated DMCs percentage.

**Supplementary Table 1.** Comparison of program features of *edgeR* and *methylKit*.

| | *edgeR* | *methylKit* |
|---|---|---|
| **Platform** | Win/Linux | Win/Linux |
| **Source** | Bioconductor, R package (v3.32.1) | Bioconductor, R package (v1.16.1) |
| **Alignment** | No | No |
| **Single/paired end** | Accepts both | Accepts both |
| **Differential methylation analysis** | Yes | Yes |
| **Concept used** | Negative binomial regression | Logistic regression |
| **Model/test** | Competitive/self-contained GLM LRT or QL F-test | Self-contained test |
| **Statistical method** | Exact text (analogous to Fisher's exact test) adapted for overdispersed data | Fisher's exact or logistic regressionc to calculate P-values |
| **Quality control and preprocessing** | Coverage normalization, removing low coverage | Library size normalization, removing low coverage |
| **Smoothing** | No smoothing involved | No smoothing involved |
| **Further analysis** | Annotation and visualization | Annotation and visualization |
| **Reference** | Chen et a. 2018 | Akalin et al. 2012 |
